# ULTRASOUND-ASSISTED MAGNETIC NANOPARTICLE-BASED GENE DELIVERY

**DOI:** 10.1101/2020.03.31.018440

**Authors:** Wei Zhang, Gaser N. Abdelrasoul, Oleksandra Savchenko, Abdalla Abdrabou, Zhixiang Wang, Jie Chen

## Abstract

Low-intensity pulsed ultrasound (LIPUS), a special type of ultrasonic stimulation, is attracting a lot of attention for both clinical and scientific research. In this paper, we report a concept of a new method using magnetic nanoparticles (MNPs) for LIPUS-assisted gene delivery. The MNPs are iron oxide superparamagnetic nanoparticles, coated with polyethyleneimine (PEI), which introduces a high positive surface charge, favorable for the binding of genetic material. Due to the paramagnetic properties of the MNPs, the application of an external magnetic field increases transfection efficiency; meanwhile, LIPUS stimulation enhances cell permeability. We found out that stimulation at the intensity of 30 mW/cm^2^ for 10 minutes yields optimal results with a minimal adverse effect on the cells. Combining the effect of the external magnetic field and LIPUS, the genetic material (GFP or Cherry Red plasmid in our case) can enter the cells. The flow cytometry results showed that by using just a magnetic field to direct the genetic material, the transfection efficiency of HEK 293 cells that were treated with our MNPs was 56.1%. Coupled with LIPUS stimulation, it increased to 61.5% or 19% higher than the positive control (Lipofectamine 2000). In addition, compared with the positive control, our method showed less toxicity. Cell viability after transfection was 63.61%, 19% higher than with the standard transfection technique. In conclusion, we designed a new gene-delivery technique that is affordable, targeted, shows low-toxicity, yet high transfection efficiency, compared to other conventional approaches.

**The Graphical Abstract:** 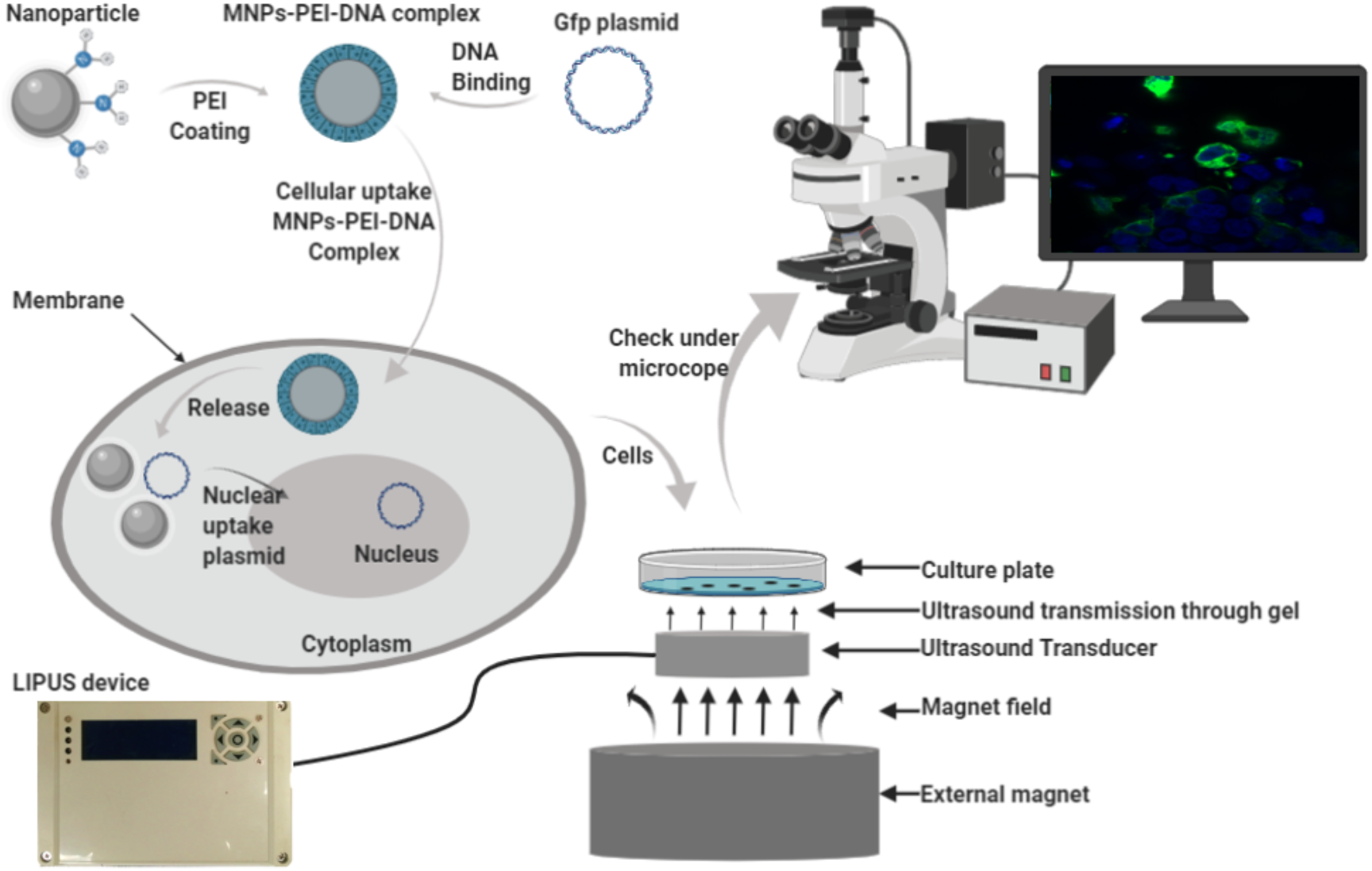

## 1. Introduction

Gene delivery is now a popular research area with high demand on the market, and applications in both clinical and scientific biomedical research [1,2]. The applications include, but are not limited to, treating cancers, immune-deficient diseases, and genetic diseases [3]. Mammalian cells have a selectively permeable plasma membrane that protects them from the external environment. Effective methods to transfect cells are needed. For the delivery of genetic material into the nucleus of the cell, two approaches can be suggested: increasing the cell membrane permeability and thus facilitating the penetration of the target gene, or developing a carrier that can go through the cell membrane, carry the gene and deliver it to the nucleus. Based on these two different pathways, gene delivery utilizes either chemical or physical methods [4,5]. The chemical approaches can be further divided into viral and non-viral approaches [4]. The ideal carrier should be low cost, with high loading capacity, high stability, no or low toxicity, and easy to use [11]. The viral-vector system approach is the most common and widely used method [3], which can achieve very high transfection efficiency. However, the safety concerns related to immunogenicity and the high cost remain the main limitations [5]. Non-viral methods include liposome-based methods [12], calcium phosphate precipitation [13], cationic polymers [14,15] (such as polyamidoamine dendrimers and PEI [16]), and nanoparticle-based hybrids [17]. The cationic liposomes are the most commonly used non-viral delivery system for gene delivery. They can reach most of the requirements of the ideal characteristics with the significant drawbacks of high toxicity and the inflammatory responses [12]. Calcium phosphate precipitation and PEI get low transfection efficiency and high cytotoxicity [13]. Nanoparticles are submicron-sized polymeric particles, due to the sub-cellular and sub-micron size range, they can penetrate tissues more efficiently [19]. MNP is one of the traditional nanoparticles and is also a popular carrier for gene delivery [18]. MNP can overcome the weaknesses of other traditional carriers, like high toxicity due to which they can only be used *in vitro* [20]. The external magnetic fields applied on the target site not only can enhance the transfection, but also target the gene to a specific site without the side effects on other tissues. Due to this, MNPs can be tunable and focus on the target area, yet they still have some drawbacks like low transfection efficiency and toxicity [31].

Besides the chemical approach, the physical delivery methods are attracting more and more research interests, including the application of the electric field [6], the acoustic method [7], and physical injection [8], to disrupt the cell membrane and let the DNA pass through it more efficiently. Some physical methods have shown sufficient delivery efficiency and can be applied to most of the cell types and are available for commercial use [9]. However, the main limitations are the cytotoxicity and the inability to be used in humans. Also, operational and equipment requirements are complicated and costly, and sometimes with low efficiency and repeatability. Acoustic methods are another physical approach to transfect the cells with the advantage of easy repeatability and good stability, yet with low transfection efficiency compared with other methods [10].

The ultrasound method is one of the acoustic transfection methods mentioned above, which is characterized by frequencies greater than 20 kHz. Ultrasound has been used for various applications, including diagnosis, surgery, and therapy for a long time [22] [23]. At its early applications, researchers focused on the treatment produced by using the thermal effects of ultrasound, while nowadays more researchers are paying attention to the non-thermal characteristic, including acoustic cavitation and mass transfer enhancement [24]. For ultrasound medical applications, the safe range of the intensity is between 0.05 W/cm^2^ and 100 W/cm^2^ [25,26]. LIPUS is a particular type of ultrasounds shown in Fig 1a, which generates at a frequency of 1-3 MHz and repeats at 1 kHz with a duty cycle of 20% to deliver a low-intensity pulsed wave [27]. It has been proven to be very safe for human use for many aspects of the medical applications, such as bone healing [28], inflammation inhibiting [29], soft-tissue regeneration [30], and the induction of cell-membrane porosity. LIPUS has also been proven to help division and proliferation in many types of cells, such as algal cell [39, 40], stem/progenitor cell [37] and mammalian cell [38] and can also increase CHO cells growth and antibody production[35], increase cell permeability [39] and enhance gene delivery by using microbubble [36]. Again, the safe operational intensity range of LIPUS is between 0.02 and 1 W/cm^2^ and treatment durations of 5 - 20 minutes per day. Because of its low intensity, LIPUS has almost no thermal effects [33].

**Figure 1:**
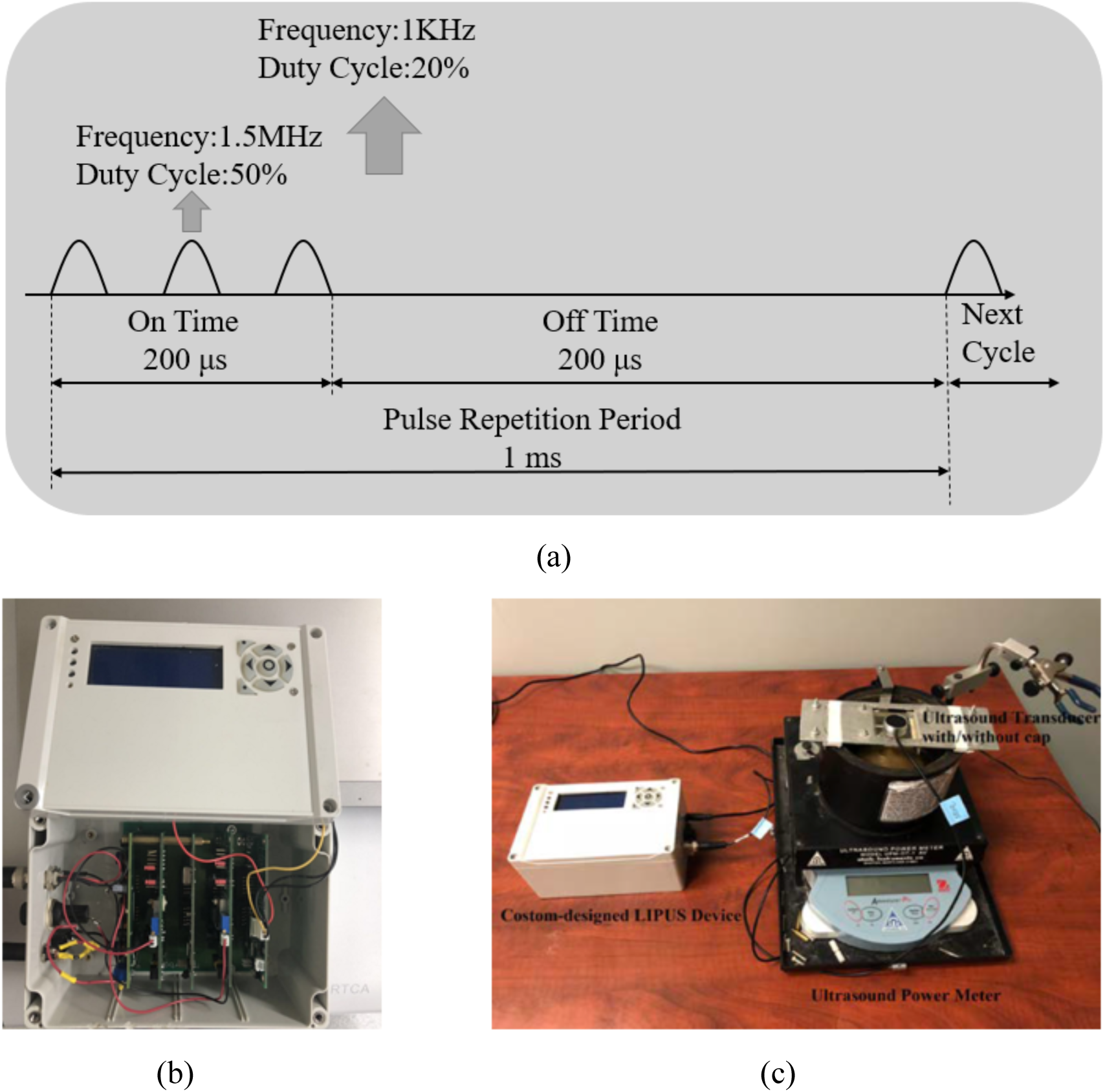
(a) A particular example of the LIPUS waveform. (b) LIPUS generation box, (c) photograph of the setup for measuring acoustic impedance.

Because LIPUS was previously shown to promote cell growth and enhance cell membrane permeability, in this study, we considered combining the LIPUS device application to enhance genetic material delivery into the cells using the PEI coated MNPs [31], [32]. Performing the cell transfection with LIPUS and MNPs under the external magnetic field, we achieved high transfection efficiency with low cytotoxicity. We present a new concept of integrating the physical and chemical gene delivery approaches by introducing LIPUS to support gene transfection using MNPs under the influence of the magnetic field. The plasmid of interest (GFP and Cherry Red plasmid) is firstly bound to the MNPs through PEI before introducing them to the cells. Then we investigate the impact of applying the magnetic field in combination with the LIPUS on the transfection of the targeted plasmid. We also examine the impact of the LIPUS on cell proliferation and viability to identify the proper ultrasound intensity and best duration that cells can tolerate. We used the fluorescent microscope to see whether this approach work or not and then employed flow cytometry to evaluate the transfection efficiency and compare the results with those obtained from using Lipofectamine 2000 as a positive control. Furthermore, we also localized the transfected genes in the targeted cells using confocal microscopy.

## 2. Materials and Methods

### 2.1 Chemicals and Materials

Anhydrous Ethylene glycol (EG) 99.8%, Ferric chloride hexahydrate (FeCl_2_.6H_2_O) ≥ 99 %, and anhydrous Sodium acetate ≥ 99% (NaAc), and branched Polyethyleneimine PEI with average molecular weight (M.W.) 25 kDa were purchased from Sigma-Aldrich and used without further purification. HEK 293T cell (human embryonic kidney cells) were purchased from ATCC, Minimum Essential Medium (MEM), Feta Calf Serum (FCS), Penicillin/Streptomycin, PBS, Lipofectamine 2000 and DAPI were purchased from Thermofisher. Zombie Aqua was purchased from Biolegend. We used Milli-Q water with the resistivity of 18.2 MΩ from the Millipore Milli-Q Advantage A10 purification system in all experiments.

#### Magnet

The magnet, made of neodymium (rare earth) with a diameter of 10 cm and a thickness of 10 cm, was purchased from Applied Magnets (Plano, Tx, USA).

### 2.2. Cell culture

HEK 293T cells were cultured in high glucose MEM medium, supplemented with 10% FCS and 1% penicillin/streptomycin at 37°C and 5% CO_2._ Cells were passed to a 12-well plate before the experiment, and transfection was performed in the 12-well plate once the cells reached 60-80%, confluency, which is recommended for the transfection.

### 2.3. Synthesis and functionalization of MNPs

MNPs were synthesized using the hydrothermal method, according to our previously reported work [20, 31]. In short, a reaction mixture containing 10 g 1,6-hexanediamide, 2.0 g FeCl_3_•6H_2_O, and 4.0 g sodium acetate trihydrate (NaAc •3H_2_O) in 50 mL of ethylene glycol (EG) was vigorously stirred at 85 °C for 2 h until a resulting transparent solution was obtained. To complete the reaction, the solution was sealed in a 100 mL- Teflon-lined stainless-steel autoclave and put the oven for 12 h at 200 °C. To collect the product, the MNPs solution was cooled down to room temperature, and MNPs were collected with the help of a magnet and further redispersed in milli-Q water by sonication for 15 min. MNPs were washed with water three times, where the MNPs were redispersed by sonication and collected each time with the help of the magnet. Then we also washed with absolute ethanol following the same method to ensure the complete removal of unreacted materials. Finally, the prepared MNPs were dispersed in 100 mL milli-Q water for characterization and further use.

For functionalization MNPs were treated with 5% glutaraldehyde solution for 2 hours, then washed three times with Milli-Q water and further coated with 1 mg/mL solution of PEI (25 KDa). PEI is known to assist in cell transfection, yet it is toxic to the cells. When combined with MNPs and due to the glutaraldehyde treatment, PEI/DNA complex with the MNPs can overcome the high cytotoxicity and instability of the PEI/DNA complex [31].

The size of nanoparticles was estimated using Hitachi HF-3300 Transmission Electronic Microscope (300 kVTEM, where we measured the size of 150 nanoparticles using ImageJ software, and the size of the nanoparticles was calculated based on the histogram of the size distribution.

The ζ potential and the hydrodynamic size of the MNPs were measured using Zetasizer Nano ZS Malvern Panalytical. The prepared MNPs were ∼24 nm in size, as shown in Fig.2a. The hydrodynamic size and zeta potential for our nanoparticles were evaluated and shown in Fig. 2b.

**Figure 2:**
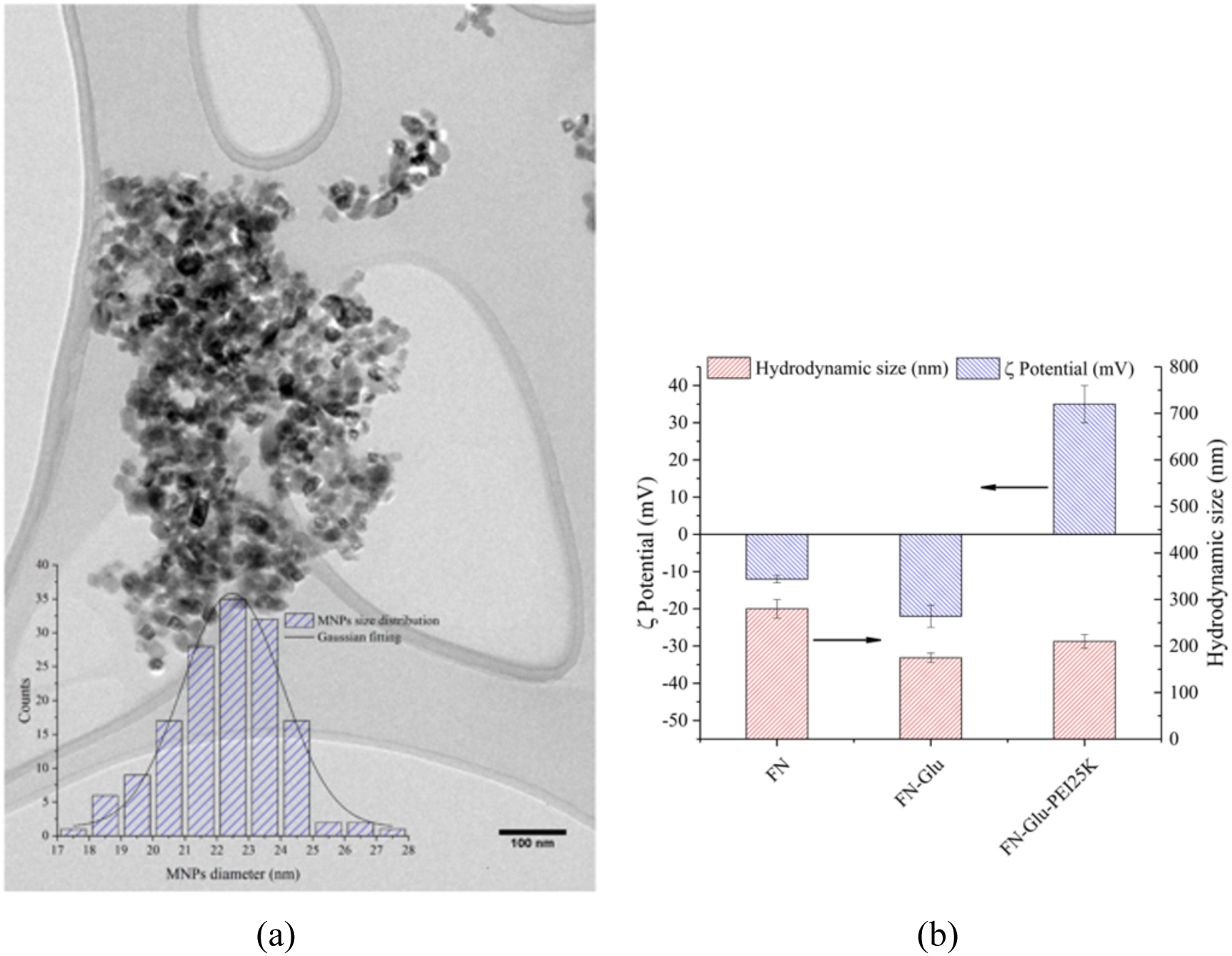
(a)MNPs size distribution under TEM. (b) Hydrodynamic size and ζ potential for particles.

### 2.4 Ultrasound stimulation device

Cells were exposed to ultrasound stimulation using the LIPUS device developed previously in our lab. The LIPUS device outputs a square wave with a frequency of 1.5 MHZ. The repetition rate is 1 kHz, and the duty cycle is 20%. We can adjust the output voltage from 1.25V to 12.5V by adjusting the potentiometer. Along with the increase of the ultrasound intensity, the output voltage increases too. The PCB board is shown in Fig.1b, and has a total of 6 boards in the box, including a motherboard, a control board, a power board, an ultrasound board, and two driver boards. The motherboard is used to connect all other boards. The control board is used to control the ultrasound intensity and duration. The ultrasound board is used to supply an ultrasound signal with a frequency of 1.5 MHZ and a repetition rate of 1 kHz, and the driver board is used to provide enough voltage and current to drive the transducers. Ultrasound settings can be controlled and adjusted.

#### Ultrasound transducers

Two ultrasound transducers we used in the LIPUS device were purchased from American Piezo Company (APC) International, Ltd (Mackey Ville, USA). The piezo-crystal 880 inside the transducer has a diameter of 25 mm and a thickness of 12.5 mm. Although the diameter of each well in a 12-well cell culture plate is 22 mm, the diameter of the transducer is slightly bigger than the well. However, it does not affect the results because we purposely calibrate the transducer with the intensity of 30mW/cm^2^, not the overall power. It has a resonant frequency of 1.5 MHz and a piezoelectric charge constant d33 of 215 m/V. The ultrasound power meter we used to measure the ultrasound intensity was purchased from Ohmic Instruments Co., Maryland, USA, and the model is UPM-DT-1AV. The diameter of the transducer is 25 mm, and thus its effective area is 4.9 cm^2^. For instance, if we want to have the output intensity of 30 mW/cm^2^, the measured output power should be 4.9×30=147 mW=0.147 W. The minimum measurement changing value of this power meter is 0.002 W. Therefore, in a real operation, we adjust the resistance of the potentiometer to get the readings as 0.146 W, or 0.148 W. Each of the transducers was calibrated before the experiment using a degassed water tank, in which transducer was fixed using a holder until the reading of the output was stable.

### 2.4. Cells Transfection

Transfection was performed at 60-80% cell confluency. GFP plasmid or Red Cherry plasmid were used as genetic material to transfect. 1ug purified plasmid was mixed with 3 µL MNPs both previously diluted in 500 µL serum-free medium to a total volume of 1000 µL and then thoroughly mixed and left for 30 min. The serum can interfere with the formation of the complex of DNA with MNPs. [41] After 30 minutes, the medium was removed from the cell wells in a 12-well plate and replaced with the mixture of MNPs-DNA complexes. To direct the MNPs into the cells, we then put the plate on the LIPUS/magnet device and treated for 10 minutes. After that, we stopped the LIPUS, and incubated the cells on the magnet for 4 more hours. The medium was then replaced with 10% serum MEM medium, and cells were further grown in the incubator for up to 48 hours.

For the Lipofectamine 2000 transfection, 1 µg purified plasmid was mixed with 3 µL lipofectamine in serum-free MEM and left for 30 mins. We added the DNA-lipofectamine complex to the cells after the 30-minute incubation.

### 2.5. Experimental Setup

After mixing cells with the genetic material of interest and transfection reagent (MNPs or Lipofectamine), the cell plate was placed on the ultrasound transducer with the ultrasound gel connection for 10 min treatment. After that, the plate was put on top of a strong magnet below the transducer. After the treatment, the ultrasound device was removed, and the cells were kept on the magnet for 4 hours. Transfection efficiency was checked within 24-48 hours after the experiment.

### 2.6. Transfection evaluation/characterization

Cell transfection efficiency was evaluated within 48 hours after the experiment using several methods. Confocal microscopy images were obtained using the Zeiss LSM 710 confocal microscope. For the confocal microscope imaging, cells were cultured and transfected on the cover slip using the same above-mentioned protocol, and after 48 hours were fixed in the 4% PFA and stained with DAPI. Fluorescent microscope images were obtained by Zeiss, Axiovert 200 fluorescent microscope. For the qualitative evaluation of transfection, fluorescent microscopy was used. Cells after transfection were checked within 24-48 hours for the fluorescent protein expression. As a negative control, untreated HEK 293T cells were used. Both groups of cells were fixed in 4% freshly prepared paraformaldehyde (PFA) for 10 minutes.

## 7. Flow Cytometry

The quantification of the transfection efficiency was performed using Attune X Flow Cytometer. Flow cytometry was performed with Zombie Aqua as our cell viability dye in dilution of 1:250, which was selected based on the pre-experimental titration and gave the clearest separation of the dead and alive cells. In the Flow Cytometry experiment, aside from experimental groups of samples, we used compensation controls, positive control, and negative control to assure the accuracy of the results.

a. For the negative control, untreated cells without any genetic materials and viability dye were used.
b. As there were two main fluorophores, we used two compensation controls to prevent the spills: For Zombie Aqua compensation, we used cells, stained with viability dye; For GFP or Cherry Red compensation, we used cells transfected with a plasmid (GFP or Cherry respectively) and Lipofectamine as transfection agent.
c. For the positive control, the purified plasmid was used for transfection, and Lipofectamine as a golden standard for transfection available on the market, and cells were stained with Zombie Aqua after 48 hours.
d. For the experimental group, two groups were set to evaluate the effect of LIPUS on the MNPs transfection and cell viability. First group: cells, plasmid, and MNPs were treated with magnet for 4 hours and stained with viability dye after 48 hours. The other group was additionally treated with LIPUS device (10 mins duration at 30mW/cm^2^ intensity) and a magnet for 4 hours and then stained after 48 hours.

## 3. Results and Discussions

In this paper, we combined both physical and chemical approaches in the development of a high-efficient gene delivery method with low-cytotoxicity. From the previous work done in our group as well as by other groups and reported in the literature, we knew that LIPUS could transiently increase cell membrane permeability [39] as well as can be beneficial for cell viability [35,38]. Our goal was to evaluate whether ultrasound can enhance the entry of genes into the cells, when used with MNPs as a transfection tool.

### 3.1 Selecting Optimal Ultrasound Condition

Because different ultrasound parameters, such as wave intensity, treatment duration and frequency of the treatment, can have a significantly different effect on the cell growth and the membrane permeability [35,39], we needed to select the optimal conditions of the LIPUS stimulation for our purpose among five different ultrasound intensities and durations (1.No LIPUS stimulation; 2. LIPUS at 30 mW/cm^2^ for 5 minutes; 3. LIPUS at 40 mW/cm^2^ for 5 minutes. 4. 30 mW/cm^2^ for 10 minutes. 5. 40 mW/cm^2^ for 10 minutes). Based on previous successful work done in our group on various types of cells, 30 mW/cm2 was selected as the optimal ultrasound condition for mammalian cells [38]. For the successful penetration of ultrasound waves to stimulate the cells, ultrasound gel has to be applied on the surface of the ultrasound transducer as ultrasound does not spread through the air. If we do not use the ultrasound gel, the transmission coefficient of sound intensity can be 0.00923%. When we use the ultrasound gel, the transmission coefficient of sound intensity can be 31.1%, and thus the ultrasound gel is necessary.

To select the optimal duration of LIPUS treatment, several conditions were evaluated, and cell counting was performed. The results are shown in Fig.3, where the cells treated for 10 minutes under 30mW/cm^2^ showed the best result. Consequently, we use the LIPUS device for 10 minutes under the intensity of 30 mW/cm^2^ in our transfection experiments. In our experiments, we used two ultrasound transducers at the same time to treat the cells in 2 wells, but our experiment design and set-up allow up to six wells to be treated at the same time.

**Figure 3:**
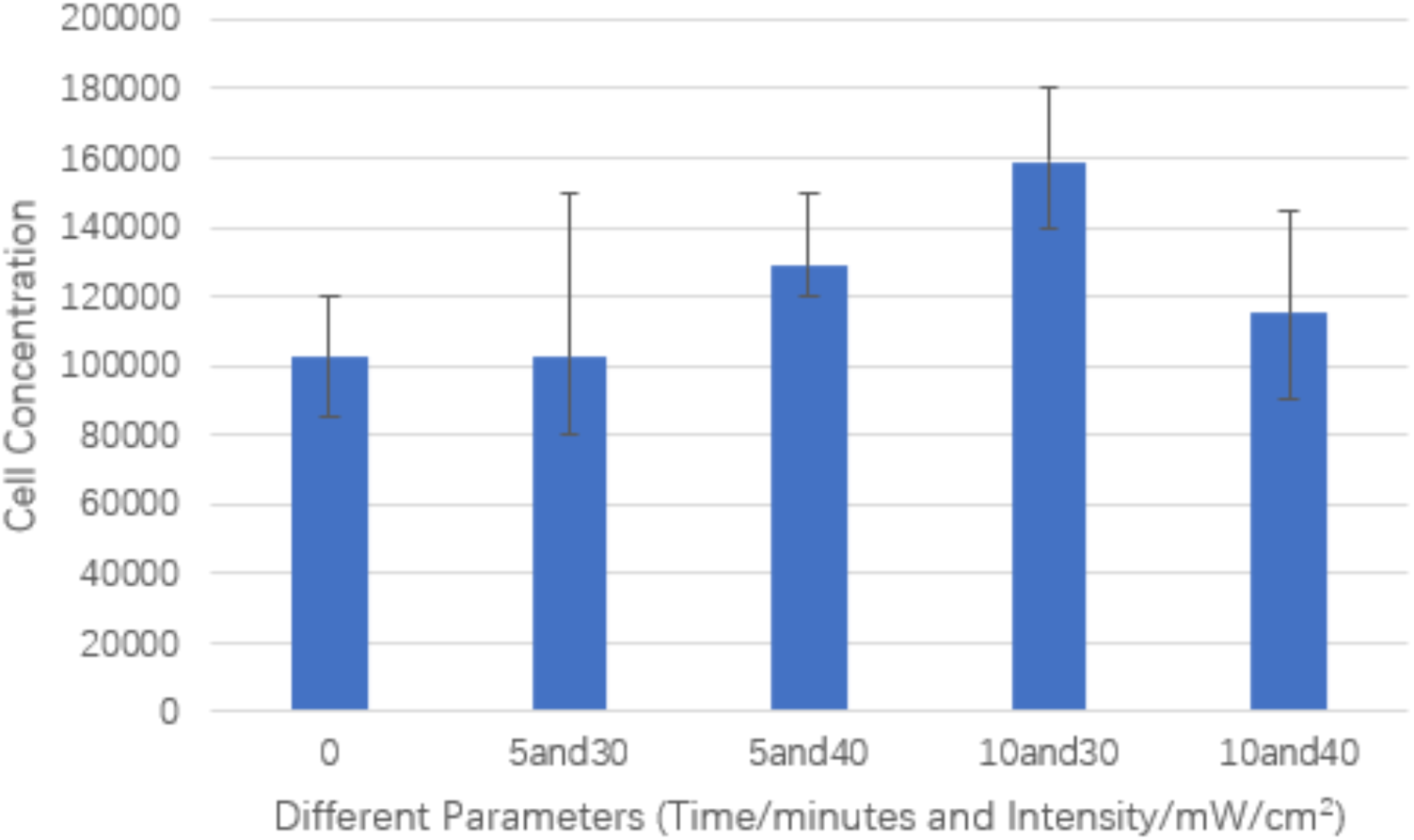
Cell proliferation after stimulation with LIPUS evaluating different intensity/duration parameters.

### 3.2 Fluorescent Microscope Results

A fluorescent microscopy allowed for an easy way to qualitatively evaluate how the combined gene delivery method worked. The negative control, which just contained the GFP plasmid in the cell plates, showed no transfected cells (Fig.4a), confirming that the plasmid itself does not cross the cell membrane without a delivery carrier. Compared to the negative control, using MNPs along with LIPUS treatment introduced the green fluorescent spots in the images, showing the cells that have been successfully transfected with the GFP plasmid and GFP expressed. These fluorescent images were good indicators that our method could work well and were very useful in the process of method development as those allowed for fast qualitative screening of each experiment.

**Figure 4:**
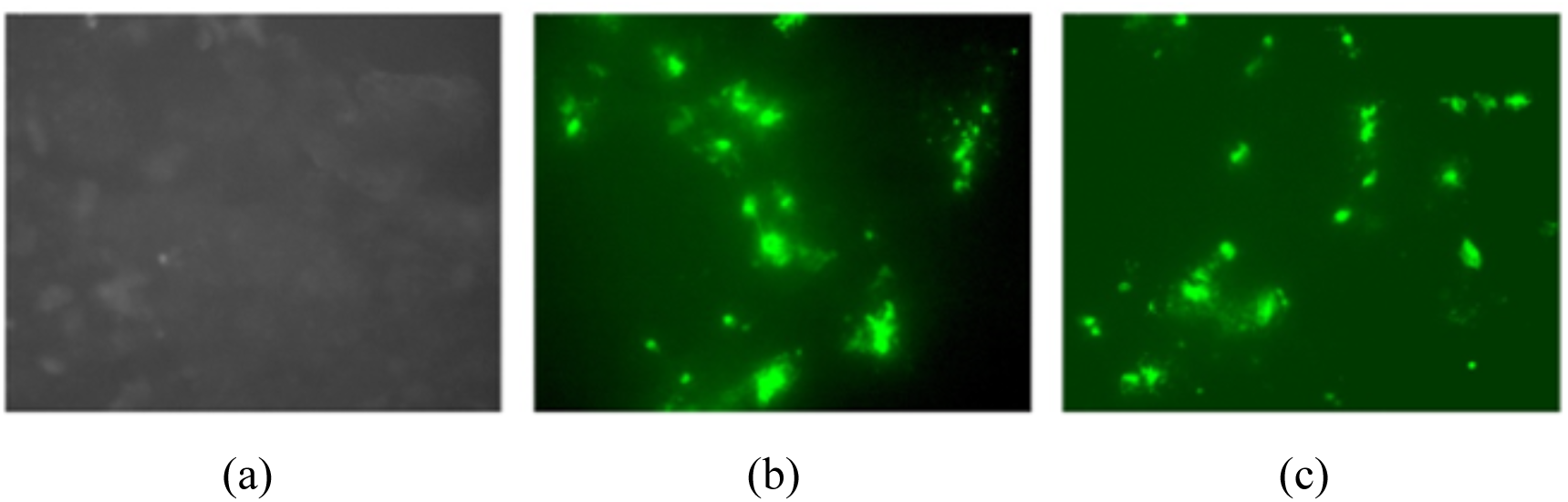
Results under Fluorescence Microscope, (a) negative control (just cells), (b) cells transfected with GFP with MNPs and (c) cells transfected with GFP with MNPs and treated with LIPUS.

### 3.3 Transfection Efficiency Using Flow Cytometry

At the same time, the fluorescent microscope images could only give us the qualitative evaluation of whether our new physical and chemical combined approach worked for gene delivery. Once we had our method developed, we had to quantitively evaluate its efficiency and whether it could be offered as an alternative to the available on the market transfection tools. We were focused on the parameters of transfection efficiency, cell viability, and how they can be compared with the results of the standard gene delivery approach using Lipofectamine 2000, a well-known and efficient transfection reagent, shown in Fig.5b. The average transfection efficiency of the Lipofectamine 2000 in our experiments was 42.62% as shown in Fig.5a, and it was within the range of normal performance of Lipofectamine 2000 working on HEK 293T cells. This result ensured that our HEK cells were always in good condition before we performed the transfection steps, and we got the proper operations during the transfection.

**Figure 5:**
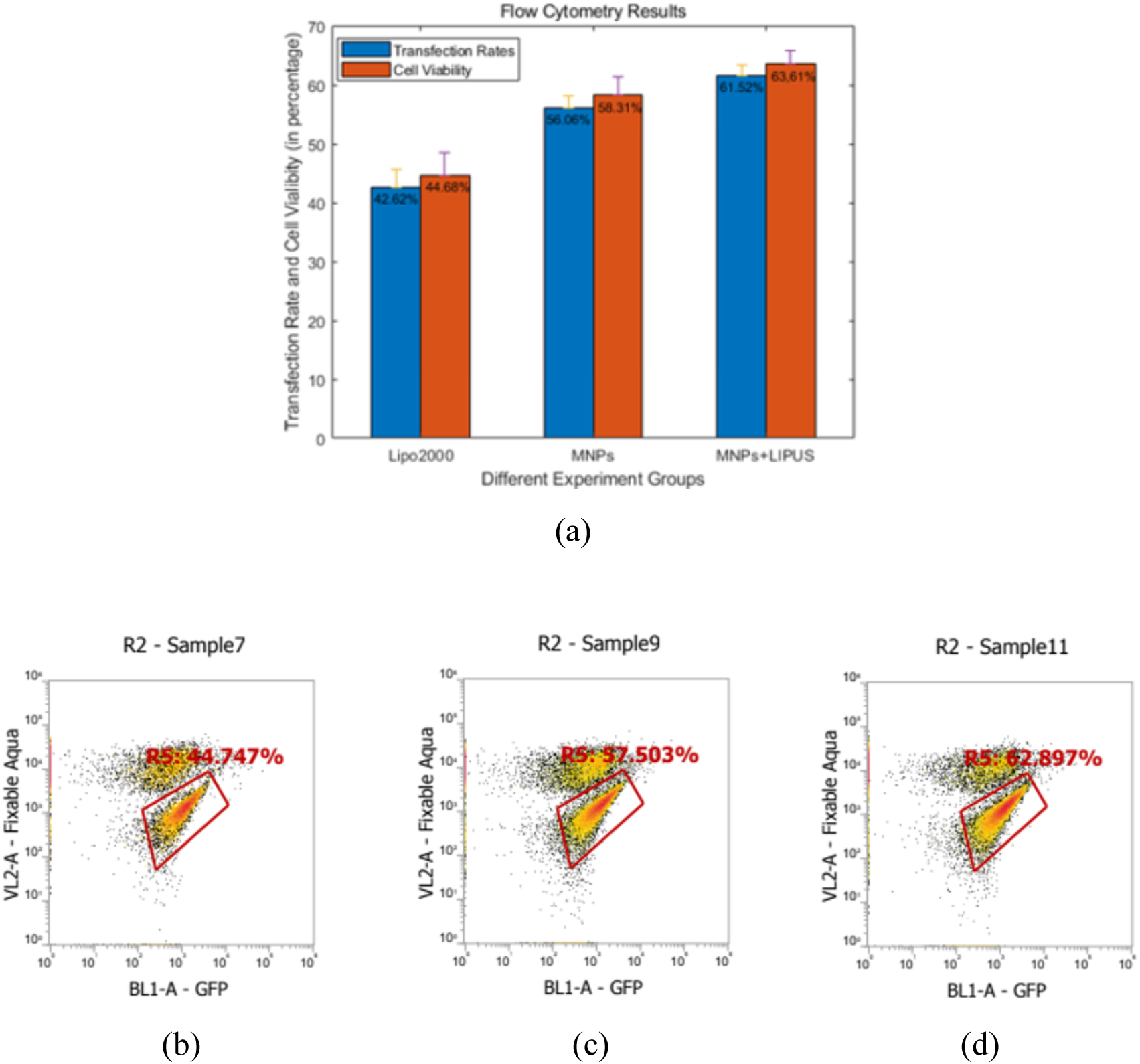
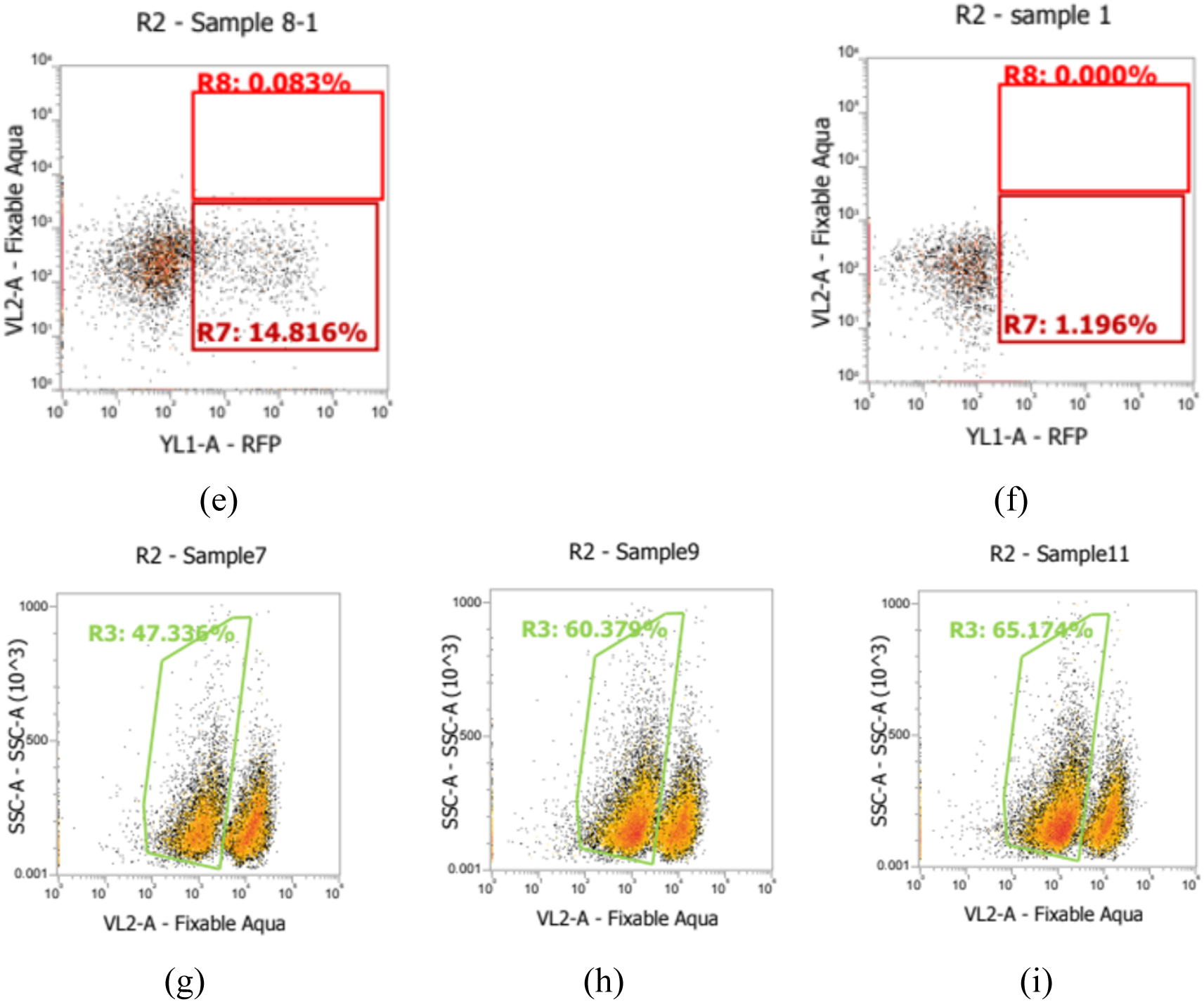
(a) Overall transfection efficiency and cell viability results. Flow cytometry histogram plots showing transfection rate using GFP and (b) lipofectamine 2000 as transfection method, (c) our MNPs and magnet, (d) our suggested method: MNPs, magnet, in combination with LIPUS treatment. (e)MNPs only, (f)ultrasound treated only. Cell viability results in the presence of Zombie Aqua viability dye when transfected with (g) lipofectamine 2000. (h) MNPs and magnet and (i) MNPs, magnet, and LIPUS.

Compared to the standard Lipofectamine 2000 reagent mentioned above, the MNPs alone and MNPs coupled with LIPUS stimulation gave us better results (shown in Fig.5a) and increased the transfection efficiency 1.3- and 1.45-fold, respectively, over the Lipofectamine 2000. Fig.5c is the histogram plot of flow cytometry, showing the GFP fluorescence of the experimental group transfected with GFP plasmid, MNPS and external magnetic field. The transfection efficiency was 57.503%, 13% higher than the Lipofectamine 2000, which indicated that the MNPs themselves could perform better than the Lipofectamine under the external magnetic field. We also ran an experiment with MNPs, but without the application of the external magnetic field and the transfection efficiency was only 14%, as shown in Fig.5e. It is consistent with the idea that MNPs as a gene carrier cannot deliver the material efficiently without the external magnet field targeting.

For the development of additional LIPUS stimulation step of our method, we first selected the ultrasound condition as 10 minutes treatment with the intensity of 30 mW/cm^2,^ and then performed the transfection using our experimental set-up with LIPUS device and got the average result of 61.52%, showing the highest transfection efficiency among our samples. As a control, cells mixed with the genetic material (plasmid of interests) and treated with ultrasound were used to evaluate the effect of just LIPUS stimulation on the transfection without the MNPs/magnetic field. In that experiment, the transfection rate was only at 1% as shown in Fig.5f, indicating that LIPUS alone could not transfect the cells without the carriers. Our results showed that, though LIPUS waves could not function as a tool for transfection itself, it could permeabilize cell membranes and, coupled with another tool (i.e., MNPs in this case), could enhance gene delivery into the cells.

### 3.4 Cell Toxicity Results

In our experiment, we used viability assay (Zombie Aqua) to evaluate the viability of HEK cells in our experiments with different gene delivery approaches. Fig.5g, 5h, 5i shows three different groups of cell viability results (Group 1: Lipofectamine 2000, Group 2: MNPs/magnetic field without LIPUS stimulation, Group 3: MNPs/magnetic field plus LIPUS), and the overall results are showing in Fig.5a. The negative control group (just untreated cells) showed viability at 91.524%, which was used as the background for all the results. Lipofectamine 2000 gave us 44.68% cell viability. MNPs showed 14% higher cell viability, compared to the Lipofectamine 2000, indicating that our MNPs had lower cytotoxicity. The addition of the LIPUS device stimulation to the MNPs delivery increased cell viability further up to 63.61% after the transfection. These results showed that the LIPUS wave could stimulate cell growth and enhance cell viability.

### 3.5 Confocal Microscope Results

For confocal microscopy, the cells were transfected using the same protocol, but they were cultured on slides instead of 12-well plate. Cells were stained with DAPI in order to evaluate the location of the gene in the transfected cells. Fig 6 shows confocal microscopy results of the cells after transfection within 48 hours. In the figure, it can be seen that both with Cherry Red plasmid and GFP plasmid HEK cells were sufficiently transfected using our suggested developed technique. Also, the merged images indicate that the gene was mostly accumulated and expressed in the nucleus where the Dapi stains, confirming the successful delivery of the genetic material to the nucleus.

**Figure 6:**
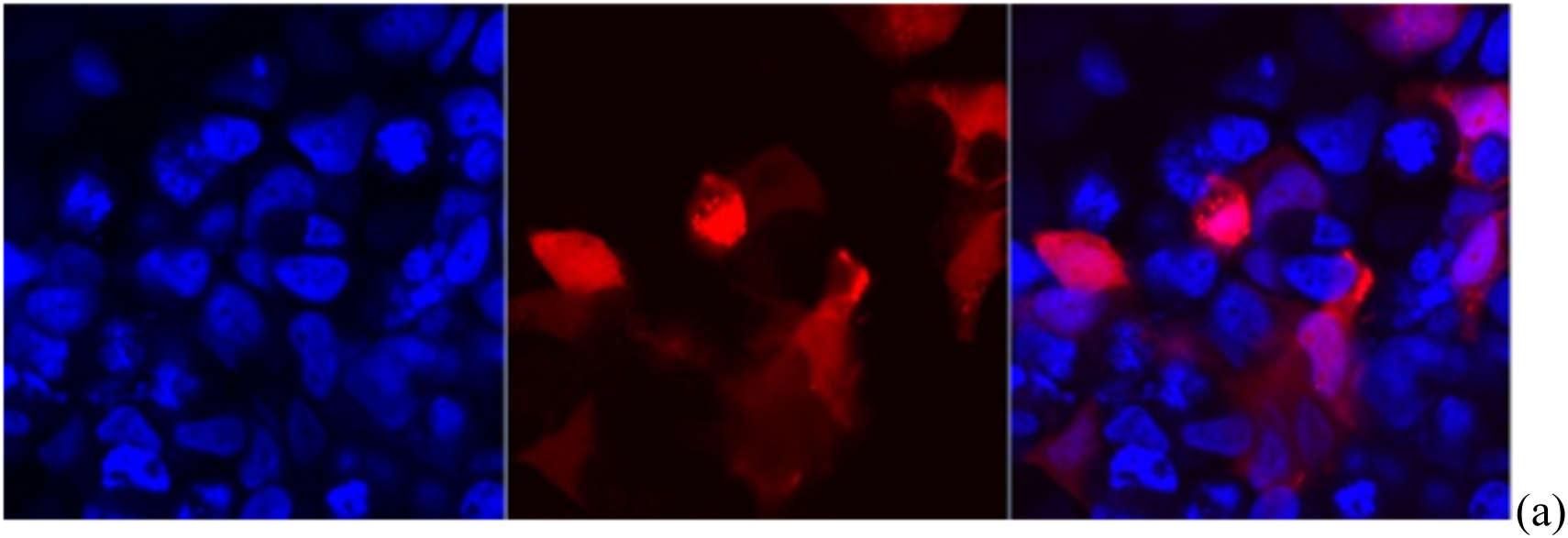
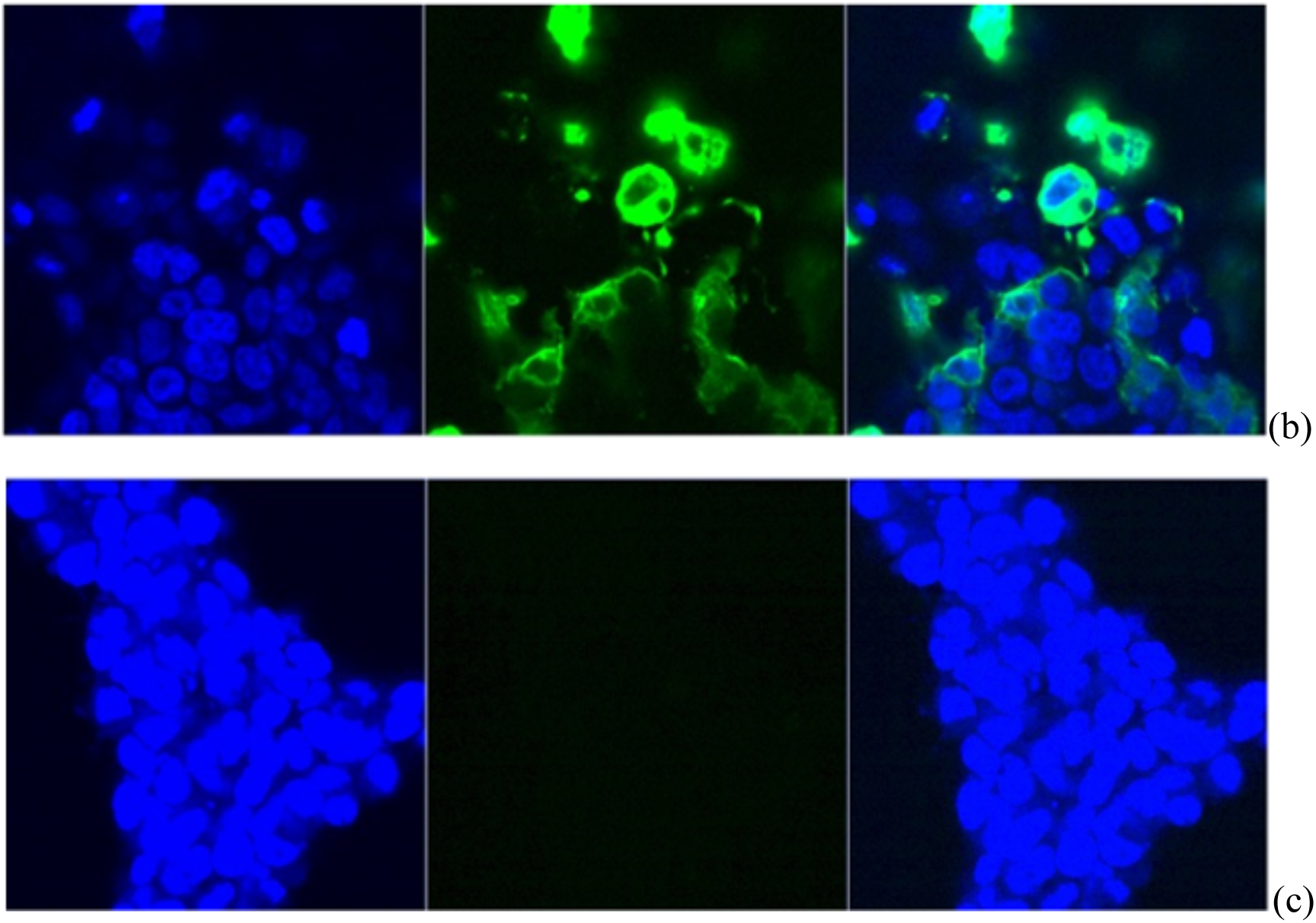
Split and merged images of cells transfected different plasmids using both MNPs and LIPUS stained with DAPI, (a) Cherry-red, (b) GFP. (c) Control group, just cells.

## 4. Conclusions

In this study, we combined application of an external magnetic field with MNPs and LIPUS stimulation for gene delivery. The uniqueness of our design is that, in addition to MNPs, used as a transfection carrier, we used the LIPUS cell stimulation to enhance gene delivery through increased cell permeability. In our experiments, we did the transfection on the HEK cells using our nanoparticles and got a 14% higher transfection rate, compared to the Lipofectamine 2000. The transfection efficiency further increases by 5%, when we add the LIPUS cell stimulation to the whole system, which was in line with our expectations. As for the cell viability, Lipofectamine is known for its cytotoxicity and showed only 44.48% cell viability in our transfection experiments. The higher percentage of cells were alive after transfection, when we used the MNPs with viability up to 58.31%. LIPUS stimulation added as an extra step during the MNPs transfection yielded even higher cell viability at 63.61%, compared to the MNPs only.

It is worth mentioning that our results and the Lipofectamine 2000 results were compared in terms of both the transfection efficiency and in cell viability, and our technique showed better performance. LIPUS was shown to promote cell permeability and let the MNPs-DNA complex pass through and thus to increase the transfection efficiency and enhance the cell viability. Because our assay is 10x cheaper than Lipofectamine 2000 and is also a chemical-based physical delivery approach, it can be an attractive gene-delivery method for other hard-transfected cells (such as prime cells and neuron cells) and *in vivo*.

## Acknowledgement

We would like to thank the funding support of MITACS. We also thank Geraldine Barron from the Cross-Cancer Institute in Edmonton, Canada for helping us with the confocal microscope and Hamid Maadi for helping us with the fluorescent microscope.

